# Molecular structures and mechanisms of DNA break processing in mouse meiosis

**DOI:** 10.1101/2019.12.17.876854

**Authors:** Shintaro Yamada, Anjali Gupta Hinch, Hisashi Kamido, Yongwei Zhang, Winfried Edelmann, Scott Keeney

**Affiliations:** Molecular Biology Program, Memorial Sloan Kettering Cancer Center, New York, NY, USA; Department of Radiation Genetics, Graduate School of Medicine, Kyoto University, 606-8501 Kyoto, Japan; Wellcome Centre for Human Genetics, University of Oxford, Oxford, UK; Department of Cell Biology and Department of Genetics, Albert Einstein College of Medicine, Bronx, NY, USA; Howard Hughes Medical Institute, Memorial Sloan Kettering Cancer Center, New York, NY, USA

## Abstract

Exonucleolytic resection, critical to repair double-strand breaks (DSBs) by recombination, is not well understood, particularly in mammalian meiosis. Here, we define structures of resected DSBs in mouse spermatocytes genome-wide at nucleotide resolution. Resection tracts averaged 1100 nucleotides, but with substantial fine-scale heterogeneity at individual hotspots. Surprisingly, EXO1 is not the major 5′→3′ exonuclease, but the DSB-responsive kinase ATM proved a key regulator of both initiation and extension of resection. In wild type, apparent intermolecular recombination intermediates clustered near to but offset from DSB positions, consistent with joint molecules with incompletely invaded 3′ ends. Finally, we provide evidence for PRDM9-dependent chromatin remodeling leading to increased accessibility at recombination sites. Our findings give insight into the mechanisms of DSB processing and repair in meiotic chromatin.

## Introduction

Nucleolytic processing of double-strand break (DSB) ends, termed resection, generates the single-stranded DNA (ssDNA) that is used for homology search and strand invasion during recombination (Symington 2014). Despite a decades-long appreciation of resection’s central role in DSB repair, however, we lack detailed understanding of resection mechanisms and the fine-scale structure of resected DNA ends in most species, including mammals. We address these issues here through genome-wide analysis of DSB resection during meiosis in mouse spermatocytes.

Meiotic recombination, which ensures homologous chromosome pairing and segregation and enhances genetic diversity, initiates with DSBs made by SPO11 via a covalent protein-DNA intermediate (**Figure 1Ai**) (Lam and Keeney 2014; Hunter 2015). Our current understanding, from the budding yeast *Saccharomyces cerevisiae*, is that endonucleolytic cleavage by Mre11–Rad50–Xrs2 (MRX) plus Sae2 nicks Spo11-bound strands, releasing Spo11 bound to a short oligonucleotide (Keeney et al. 1997; Neale et al. 2005) and providing entry points for modest 3′→5′ Mre11 exonuclease and robust 5′→3′ Exo1 exonuclease (Zakharyevich et al. 2010; Garcia et al. 2011; Keelagher et al. 2011; Cannavo and Cejka 2014; Mimitou et al. 2017). However, aside from yeast, meiotic resection mechanisms are unknown. In mouse, for example, we do not even know if EXO1 is required, and we have only a low-resolution population-average view of resection lengths that was deduced indirectly from sequencing of ssDNA bound by the strand exchange protein DMC1 (Lange et al. 2016).

**Figure 1.**
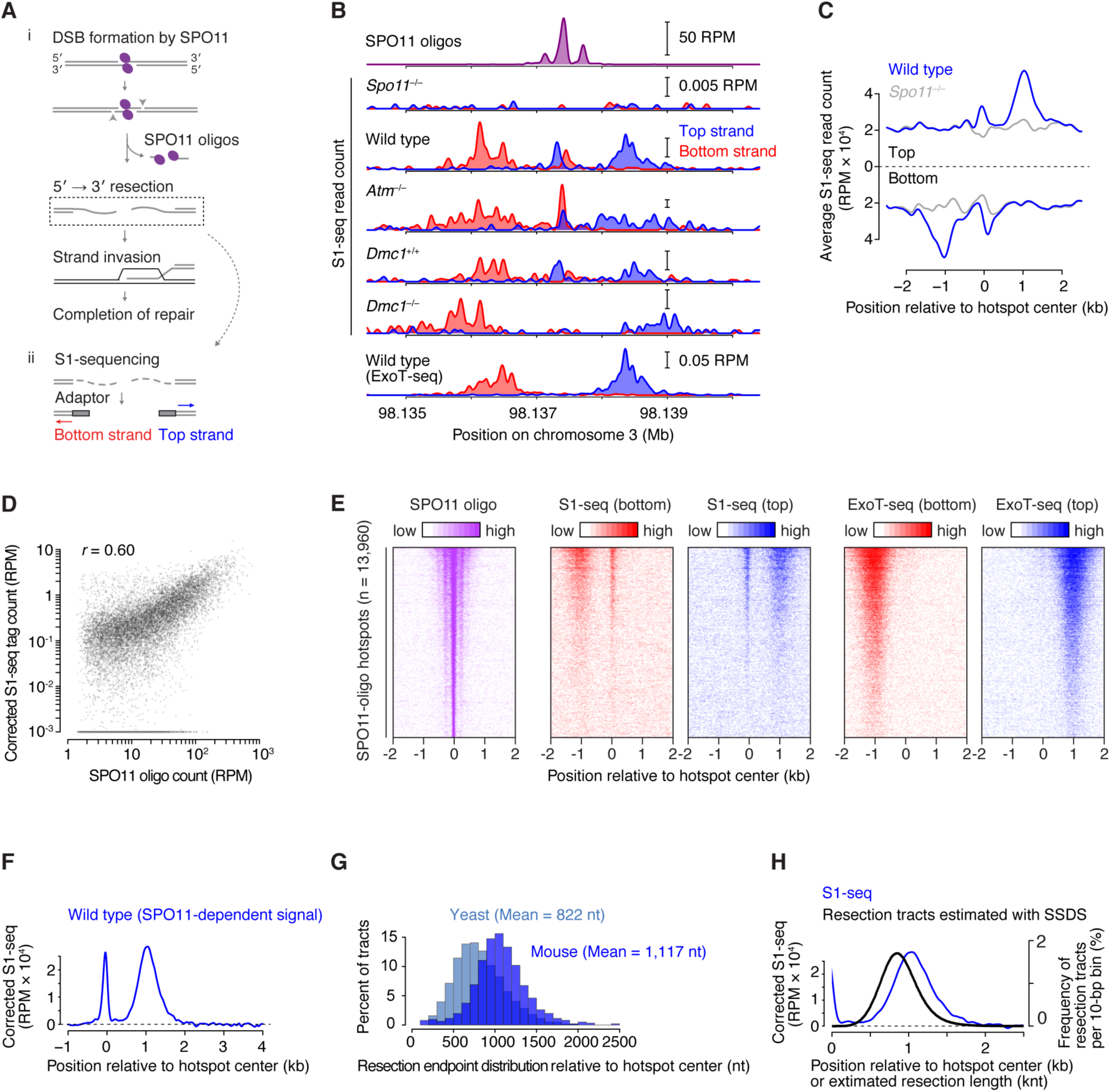
Nucleotide-resolution maps of meiotic DSB resection in mice. (A) Early recombination steps and S1-seq. (i) SPO11 (magenta ellipses) cuts DNA via a covalent protein-DNA intermediate. SPO11-bound strands are nicked (arrowheads) by MRE11 and associated factors, providing an entry point(s) for exonucleolytic resection and release of SPO11-oligo complexes. Resected ends have 3′-ssDNA ends which serve as substrates for strand-exchange proteins DMC1 and RAD51, which search for a homologous duplex and carry out strand invasion. (ii) In S1-seq, sequencing adaptors are linked to duplex ends generated by removal of ssDNA tails using S1 nuclease. (B) Strand-specific S1-seq (reads per million mapped reads (RPM)) at a representative DSB hotspot. *Dmc1*^+/+^ is a *Dmc1*-proficient control from the same breeding colony as *Dmc1*^−/–^ null on a mixed background. ExoT-seq uses an exonuclease instead of S1 endonuclease to remove ssDNA tails. SPO11-oligo sequencing data here and throughout are from (Lange et al. 2016). (C) S1-seq reads averaged around 13,960 SPO11-oligo hotspot centers in wild type and *Spo11*^*–/–*^. (D) Correlation (Pearson’s *r*) of S1-seq read count with DSB intensity measured by SPO11-oligo sequencing. Each point is a SPO11-oligo hotspot, with S1-seq signal summed from –2,000 to –250 bp (bottom strand) and +250 to +2,000 bp (top strand) around hotspot centers. S1-seq signal was background-corrected by subtracting *Spo11*^*–/–*^ signal from the wild-type signal. Hotspots with ≤0 corrected S1-seq tag counts (n = 4,050) were excluded for Pearson’s *r* calculation. Hotspots with ≤10^−3^ corrected S1-seq tag counts (n = 4,071) were set as 10^−3^ for plotting purposes. (E) Stereotyped distribution of resection endpoints around DSB hotspots. Heat maps (data in 40-bp bins) show SPO11 oligos and strand-specific S1-seq or ExoT-seq reads around DSB hotspots. Each line is a hotspot, ranked from strongest at the top. Sequencing signals were locally normalized to have the same total value at each hotspot, so that spatial patterns can be compared between hotspots of different strengths. (F) Global average patterns of SPO11-specific S1-seq reads at hotspots. S1-seq signal was background-corrected by subtracting *Spo11*^*–/–*^ signal from the wild-type signal. Top- and bottom-strand reads were then co-oriented and averaged. (G) Comparison of yeast (Mimitou et al. 2017) and mouse resection tract lengths. (H) Resection lengths measured by S1-seq are longer than those estimated using anti-DMC1 ChIP (SSDS) (Lange et al. 2016). Signal was smoothed with a 151-bp Hann filter in **B, F**, and **H**, and with a 401-bp Hann filter in **C**.

One advantage of using meiotic recombination as a paradigmatic context to study resection is that SPO11 generates numerous DSBs in a regulated fashion at a defined stage in prophase I after DNA replication (Padmore et al. 1991; Mahadevaiah et al. 2001; Murakami and Keeney 2014). Moreover, most DSBs form within narrow genomic segments called hotspots (typically <150 bp wide), which means that most DSB resection tracts emanate from relatively defined locations, facilitating their structural analysis (Lange et al. 2016; Mimitou et al. 2017).

A key determinant of hotspot locations in mice and humans is PRDM9, which contains a sequence-specific zinc-finger DNA binding domain and a PR/SET domain that methylates histone H3 on lysines 4 and 36 (Baudat et al. 2010; Myers et al. 2010; Brick et al. 2012; Wu et al. 2013). This histone methylation is required for nearby DNA cleavage (Diagouraga et al. 2018), although it remains unclear how SPO11 targeting occurs. Interestingly, PRDM9 action on the unbroken recombination partner appears to facilitate strand exchange and recombination (Davies et al. 2016; Hinch et al. 2019; Li et al. 2019), but how this is accomplished is also unknown.

In this study, we mapped the endpoints of mouse meiotic resection tracts genome-wide and at single-nucleotide resolution. We uncover previously undocumented characteristics of resection, including global patterns, locus-to-locus variation, and the genetic pathways that control resection. Our genomic sequencing method (S1-seq) also detects structures that appear to be intermolecular recombination intermediates. Unexpected properties of these intermediates in mice shed light on the mechanism of meiotic recombination and the structure of chromatin at sites undergoing recombination.

## Results

### A nucleotide-resolution map of meiotic DSB resection endpoints

To directly survey the fine-scale molecular structure of resected DSBs, we adapted the S1-seq method (Mimitou et al. 2017; Mimitou and Keeney 2018) to mouse spermatocytes (**Figures 1Aii and S1A**). Testicular cell suspensions were embedded in agarose to protect DNA from shearing, then DNA liberated by treatment with SDS and proteinase K was digested with ssDNA-specific S1 nuclease to remove ssDNA tails at DSB ends, making them blunt-ended at resection endpoints. The resulting DNA was ligated to biotinylated adaptors, fragmented by sonication, affinity purified using streptavidin, and ligated to separate adaptors at the opposite ends prior to amplification and deep sequencing. To minimize background and improve signal:noise ratio, we used testis samples from juvenile mice during the first semi-synchronous wave of spermatogenesis (12–16 days post partum (dpp)); at these ages, postmeiotic cells have not yet formed and meiotic cells are at stages when DSBs are present (leptonema, zygonema, and early pachynema) (Bellve et al. 1977; Zelazowski et al. 2017).

True resection endpoints should yield sequencing reads of defined polarity: top-strand reads for resection tracts moving away from the DSB toward the right; bottom-strand reads for leftward resection (**Figure 1Aii**). As predicted, S1-seq reads of the correct polarity were enriched adjacent to DSB hotspots for wild-type C57BL/6J (B6) mice relative to congenic mice lacking SPO11 (**Figures 1B, 1C and S1B**). Read depth near hotspots showed good reproducibility (Pearson’s *r* = 0.76 for biological replicates; **Figure S1C**) and was correlated with local DSB activity as measured by sequencing of SPO11 oligos or DMC1-bound ssDNA (SSDS) (**Figures 1D and S1D**). No similar enrichment was observed around sites that can be targeted by versions of PRDM9 that are not present in these experimental mice (e.g., PRDM9 from *Mus musculus castaneus* (CAST/EiJ) or PRDM9 carrying a humanized Zn-finger array (Davies et al. 2016); **Figure S1E**).

We conclude that S1-seq quantitatively captures *bona fide* resection endpoints. We also observed a prominent signal near hotspot centers with a polarity inconsistent with resection endpoints but consistent with expectation for S1-sensitive, intermolecular strand-exchange intermediates (e.g., **Figure 1B**). We revisit this DNA species in detail below. Because S1-seq faithfully captures DNA ends at single-nucleotide resolution in yeast (Mimitou et al. 2017; Mimitou and Keeney 2018), we assume the same is true for mouse.

### Resection is locally heterogeneous within globally stereotyped constraints

DSBs are highly clustered within hotspots in mice (**Figure 1E**, left panel), confined mostly to the relatively nucleosome-depleted regions (NDR) enclosing the PRDM9 binding sites and, to a lesser degree, the linkers between the flanking methylated nucleosomes (Lange et al. 2016). Analogous to, but more dispersed than, this clustering of DSB positions, resection endpoints showed a stereotyped distribution that was broadly similar across both sides of essentially all hotspots, with most endpoints falling within zones 0.3 to 2 kb from hotspot centers (**Figure 1E**). We obtained similar resection endpoint distributions if we removed DSB 3′ ends with the ssDNA-specific 3′5′ exonuclease ExoT (Canela et al. 2019) instead of S1 nuclease (**Figures 1B, 1E, S1B, and S1F**).

Within these broader zones of resection endpoints, however, individual hotspots showed substantial heterogeneity, i.e., reproducible peaks and valleys in the S1-seq maps (**Figures 1B and S1B**). This pattern was not from biases of S1 nuclease because peak positions appeared similar with ExoT (**Figures 1B and S1B**). Similar local heterogeneity in yeast reflects the underlying chromatin structure (Mimitou et al. 2017). We observed no clear relation of resection endpoints with preferred positions of nucleosomes, but a lack of correlation may be uninformative if nucleosome positions are variable between chromosomes in the population (**Figure S1G**).

Whatever the cause of the heterogeneity, we interpret that individual genomic locations have preferred regions for resection termination within broader zones that are defined by globally stereotyped minimal and maximal resection lengths. Previously, it was not possible to evaluate resection patterns at individual mouse hotspots because of limitations of available data (Lange et al. 2016). Our findings demonstrate the stochastic, probabilistic nature of resection termination and reveal that there is substantial variation from DSB to DSB within a cell, and even at the same hotspot between cells.

To precisely define global resection patterns, we generated a genome-wide average profile by subtracting the signal obtained in the *Spo11*^*–/–*^ mutant and then averaging top- and bottom-strand reads after co-orienting them around hotspot centers (**Figure 1F**). Alternative background-subtraction and normalization methods gave similar results (**Figure S1H**). Resection tracts averaged ∼1,100 nucleotides (nt) but with a wide, positively skewed distribution (**Figure 1G**). Of all DSB ends, 99% were resected at least 280 nt and only 1% were resected more than 1800 nt. This extent of resection predicts 440 to 660 kb of ssDNA per meiosis assuming 200–300 DSBs.

Mouse resection is more than one-third longer on average than in yeast (822 nt), although shapes of the length distributions are similar between the species (**Figure 1G**). More importantly, resection is longer by ∼200 nt that our previous estimate based on coverage from sequencing of ssDNA bound by DMC1 (SSDS) (Lange et al. 2016) (**Figure 1H**). The difference between SSDS-based imputation and our direct measurement by S1-seq may reflect technical aspects of the SSDS method (for example, partial sequence coverage of ssDNA because of the use of a foldback annealing step (Khil et al. 2012; Lange et al. 2016)). However, it is also likely that DMC1 coats only a portion of the ssDNA that usually does not include the break-distal segment closest to the ssDNA–dsDNA junction. This interpretation agrees with direct analysis of relative distributions of DMC1 and RAD51 by ChIP-seq and immunocytology (Hinch et al. manuscript submitted).

### Recombination intermediates close to DSB ends

We observed a prominent collection of S1-seq reads close to DSB hotspot centers in addition to the more distal reads from resection endpoints (**Figures 1B, 1E, 1F, and S1B**). The hotspot-proximal reads were narrowly distributed across a region shifted to the side of DSB positions as defined by SPO11-oligo sequencing (**Figure 2A**; modal shift of 35 bp). Because of this shift, we can definitively conclude that these reads are not principally from unresected DSBs, which S1-seq readily detects in yeast (Mimitou et al. 2017; Mimitou and Keeney 2018). Furthermore, the direction of the shift means that these reads have the wrong polarity to be resection tracts emanating from the DSB hotspots. These central reads made up 19.3% of the total signal around hotspots in B6 mice.

**Figure 2.**
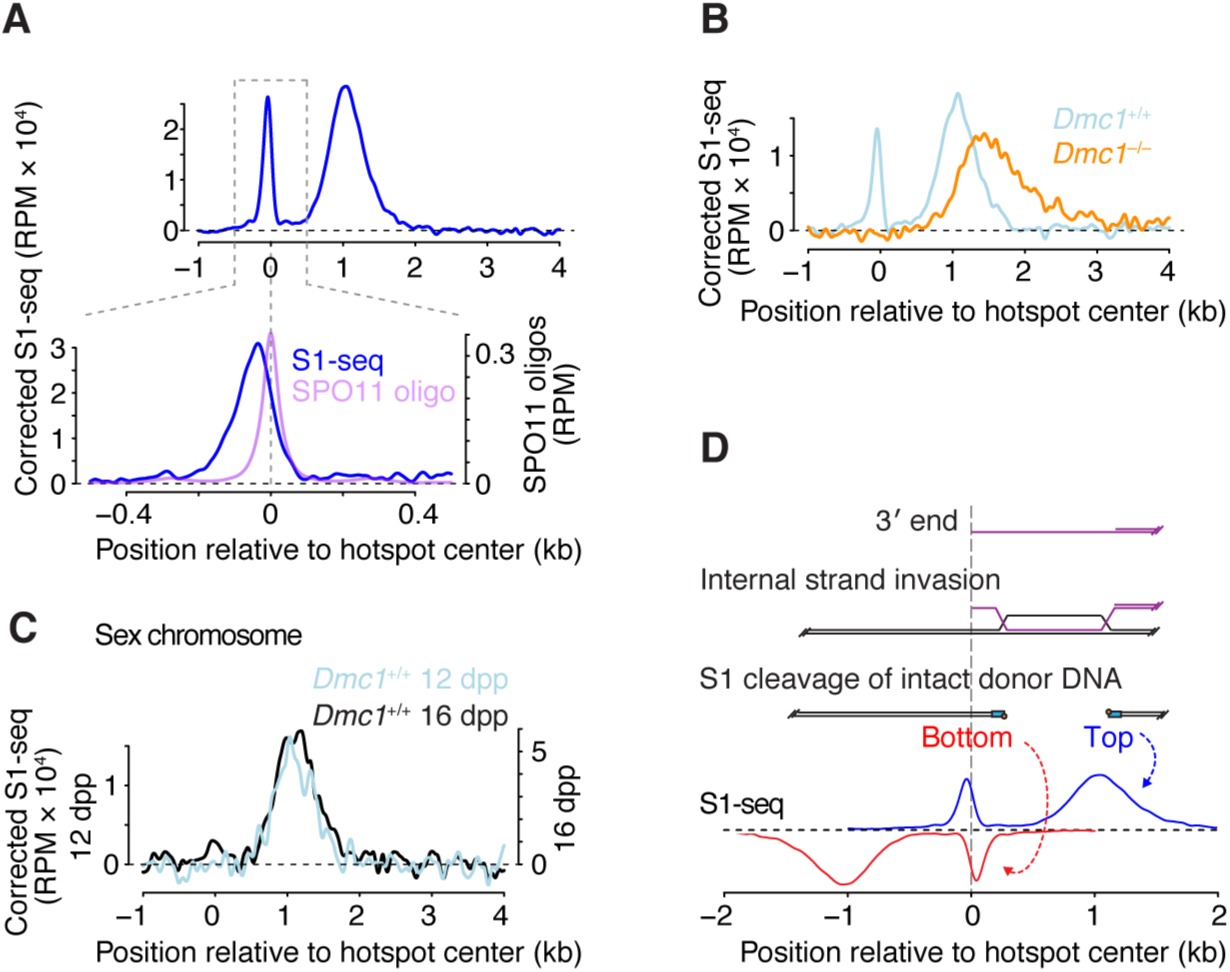
Intermolecular recombination intermediates. (A) Spatial disposition of central (hotspot-proximal) S1-seq signal. Upper panel shows the background-subtracted S1-seq signal (from Figure 1F) and lower panel shows a zoom into the region immediately surrounding hotspot centers, showing the offset of the S1-seq signal from the SPO11-oligo distribution. (B) Lack of central S1-seq signal and hyperresection of DSB ends in the absence of DMC1. Wild-type and *Dmc1*^*–/–*^ maps were prepared from animals from the same mixed background breeding colony. (C) Absence of central S1-seq signal at hotspots (n = 571) on sex chromosomes. DSB sites in the pseudoautosomal region are not included because these are too broadly distributed to be able to define hotspot-proximal vs. -distal locations. (D) Schematic illustrating how S1 cleavage of D-loops could explain the central signal. The vertical dashed line aligns the different elements of the cartoon by the 3′ end of the DSB ssDNA tail. S1 cleavage of the intact recombination partner at the DSB-proximal and -distal boundaries of the joint molecules would be expected to yield sequencing reads mapping to the bottom and top strands, respectively. See also **Figure S2C** and its legend for more details. Signal was smoothed with a 151-bp Hann filter in **A** (upper panel), **B**, and **C**, and with a 51-bp Hann filter in **A** (lower panel).

We reasoned that this central signal might be analogous to a “wrong-polarity” signal in yeast that was proposed to come from recombination intermediates of as-yet undefined structure (Mimitou et al. 2017). To test this hypothesis, we examined physical properties and genetic dependencies for formation of this signal.

The central signal was not from other hotspots nearby (**Figure S2A**), was correlated with hotspot heat (**Figure S2B**), was specific for sites being actively targeted by PRDM9 (**Figure S1E**), and was SPO11-dependent (**Figures 1B, 1C, 1F, and S1B**). We therefore conclude that the central signal requires meiotic DSB formation.

The central signal was absent in *Dmc1*^*–/–*^ mutants (**Figure 2B**), which lack a strand exchange protein essential for meiotic recombination (Bishop et al. 1992; Pittman et al. 1998). Remarkably, the signal also appeared specific for autosomes: little or no central S1-seq signal was apparent at hotspots on the X and Y chromosomes (**Figure 2C**). DSBs on non-homologous portions of the sex chromosomes persist longer than on autosomes, presumably because of a temporary barrier to using the sister chromatid as the recombination partner (Moens et al. 1997; Mahadevaiah et al. 2001; Lange et al. 2016). The absence of the central S1-seq signal at sex chromosome hotspots did not appear to be a consequence of this delayed repair because S1-seq maps made from older juveniles also had little if any of the central signal at these hotspots (**Figure 2C**). Dependence on DMC1 and absence at sex chromosome hotspots suggests that detection of the central S1-seq signal requires the ability to carry out inter-homolog recombination.

A straightforward hypothesis is that we are detecting intermolecular recombination intermediates such as D-loops (**Figures 2D and S2C**). In this model, sequencing reads are from the intact recombination partner that is rendered S1-sensitive when it is invaded by the end of the broken chromosome. If so, the central reads would be from the DSB-proximal ends of D-loops, with bottom strand reads arising from invasion by the right side of the DSB (i.e., the opposite polarity from resection endpoint reads). Sequencing reads that might come from the more distal ends of the D-loops would be expected to have the same polarity as resection endpoint reads, and might thus be masked by the larger number of resection reads (**Figures 2D and S2C.i–iii**).

One prediction from this hypothesis is that the central signal should require digestion with S1 or a similar endonuclease, i.e., it should be absent if sequencing libraries are prepared using an exonuclease (**Figure S2D**). Indeed, as predicted, maps generated using ExoT (which requires a free DNA end) failed to display the central signal (**Figures 1B, 1E, S1B, S1F, and S2D**). We note that the D-loop proposed in the canonical recombination model, in which the 3′ end of the ssDNA is fully invaded, would predict sequencing reads that line up with DSB positions (**Figure S2C.ii**). The shift that we observed instead can be simply explained if the 3′ end is not fully invaded (**Figures 2D, S2C.i, and S2C.iii**). Further implications and additional features of this model are addressed in Discussion.

We attempted to test if the central signal reflects invasion of the homolog by generating S1-seq maps from F1 hybrid mice and examining hotspots where PRDM9 targets only one of the two homologs (**Figure S2E**). The results suggest that at least some of these recombination intermediates may involve invasion of the sister chromatid. Although we could not detect the predicted signature of interhomolog joint molecules, we cannot exclude the possibility that interhomolog recombination intermediates contribute to the central signal in B6 mice (see legend to **Figure S2E**).

### Mechanism of resection

To identify molecular pathways responsible for resection, we applied S1-seq to testis samples from mutant mice. To test the contribution of EXO1, we examined mice homozygous for a knock-in mutation changing an active site aspartate to alanine (D173A, hereafter *Exo1*^*DA*^), which eliminates nuclease activity (Lee et al. 2002; Zhao et al. 2018). In yeast, the equivalent *exo1* mutation reduces resection tracts to less than one-third the normal length (Mimitou et al. 2017). Surprisingly, however, *Exo1*^*DA*^ mice showed only a modest (∼10%) decrease in resection length (**Figure 3A**). Thus, EXO1 is not a major participant in 5′→3′ resection in mouse, or it is substantially redundant with one or more other resection activities.

**Figure 3.**
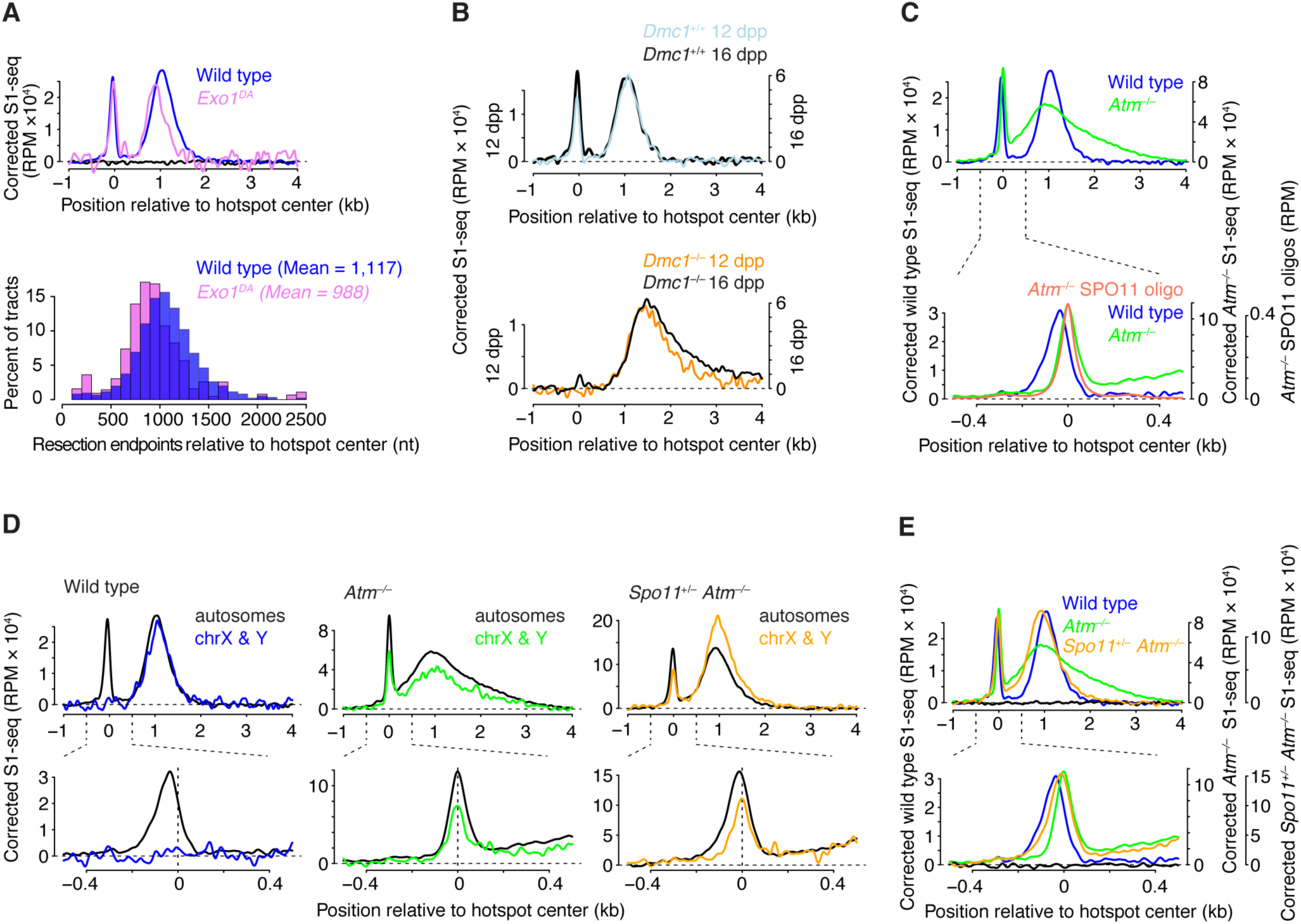
Genetic control of meiotic resection. (A) EXO1 exonuclease activity contributes only modestly to the full extent of resection length. (B) Hyperresection is greater in samples from older *Dmc1*^*–/–*^ animals (16 dpp vs. 12 dpp), which are enriched for spermatocytes at later stages in prophase I. (C) Altered resection in *Atm*^*–/–*^ spermatocytes. The lower panel shows a zoom into the region around hotspot centers, showing coincidence of the central S1-seq signal with SPO11 oligos in *Atm*^*–/–*^, unlike in wild type. (D) Unlike in wild type, the central signal in ATM-deficient mutants (with or without *Spo11* heterozygosity) is prominent at hotspots on sex chromosomes. (E) *Spo11* heterozygosity modifies the resection defects caused by ATM deficiency. Signal was smoothed with a 151-bp Hann filter in **A, B**, and **C**-**E** (upper panel), and with a 51-bp Hann filter in **C**-**E** (lower panel).

In *Dmc1*^*–/–*^, DSBs were resected further than normal (**Figures 1B, 2B, and S1B**). Moreover, more of the very long resection tracts were present in testes from older mice (16 dpp versus 12 dpp), whereas resection tract lengths in wild type were similar across these ages (**Figure 3B**). These results reveal that mice lacking DMC1 hyperresect their DSBs, and suggest that hyperresection continues progressively. Both properties are reminiscent of *dmc1* mutant yeast (Bishop et al. 1992; Mimitou et al. 2017).

The DSB-responsive kinase ATM (ataxia telangiectasia mutated) is a key regulator of DSB formation, apparently controlling a negative feedback circuit whereby DSBs inhibit formation of other DSBs nearby on the same chromatid or its sister (Lange et al. 2011; Garcia et al. 2015; Lukaszewicz et al. 2018). The ATM ortholog in yeast (Tel1) also promotes meiotic resection (Joshi et al. 2015; Mimitou et al. 2017), so we asked if this function is conserved.

S1-seq maps from *Atm*^*–/–*^ mice revealed drastic changes (**Figures 1B, 3C, and S1B**). Resection endpoints were spread over a much wider area, with many tracts shorter than in wild type and many tracts longer (**Figure 3C, upper panel**). This unexpected mix of hypo- and hyper-resection indicates that ATM controls, directly or indirectly, the extent of resection.

A central S1-seq signal was present in *Atm*^*–/–*^ mice, but with an abnormal spatial disposition that was highly coincident with DSBs (**Figure 3C, lower panel**). Unlike in wild type, the central signal in *Atm*^*–/–*^ was readily apparent at hotspots on the sex chromosomes (**Figure 3D**). These findings suggest that *Atm*^*–/–*^ mutants accumulate unresected DSBs (as in *tel1* yeast (Joshi et al. 2015; Mimitou et al. 2017)) and do not accumulate intermolecular recombination intermediates. The central signal was 14% of the total S1-seq signal around hotspots in *Atm*^*–/–*^.

We sought to determine which changes are more likely to reflect direct requirements for ATM by performing S1-seq in *Spo11*^*+/–*^ *Atm*^*–/–*^ mice. *Atm*^*–/–*^ single mutants experience a strong increase in DSB formation, estimated at ≥ 10 fold based on quantification of SPO11-oligo complexes (Lange et al. 2011). Their spermatocytes also experience severe recombination defects and catastrophic meiotic failure (Barlow et al. 1996; Barchi et al. 2005). However, reducing *Spo11* gene dosage attenuates the increased DSB numbers (Lange et al. 2011) and suppresses many of the meiotic defects of *Atm*^*–/–*^ mutants (Bellani et al. 2005; Barchi et al. 2008), suggesting that at least some of the recombination problems in ATM-deficient spermatocytes are a secondary consequence of the massively increased DSB load rather than reflecting direct roles of ATM per se.

S1-seq patterns differed considerably in *Spo11*^*+/–*^ *Atm*^*–/–*^ mice compared with *Atm*^*–/–*^ single mutants or wild type (**Figure 3E**). Hyper-resection was essentially eliminated, but a subset of DSBs remained under-resected, giving an average resection length of 978 nt (88% of the average in wild type) (**Figure 3E**, upper panel). The distribution of the central signal also changed, giving an intermediate pattern between wild type and *Atm*^*–/–*^, which we interpret as a mixture of recombination intermediates and unresected DSBs (**Figure 3E**, lower panel). This interpretation is supported by the retention of a DSB-coincident central signal at sex chromosome hotspots (**Figure 3D**).

Because reducing *Spo11* gene dosage fully suppressed hyperresection and partially suppressed the defect in forming intermolecular joint molecules, we conclude that excessive exonucleolytic processing and apparently defective strand exchange in *Atm*^*–/–*^ mutants are indirect consequences of a high DSB burden when ATM is absent. In contrast, the retention of a population of unresected DSBs and the decrease in resection tract length suggest that these defects are more directly tied to ATM function per se. We therefore conclude that ATM is critical for both the initiation and extension of meiotic DSB resection in mice. Strikingly, these aspects of the *Atm*^*–/–*^ mouse phenotype are the ones most reminiscent of *tel1* mutant yeast (Mimitou et al. 2017), suggesting wide conservation of ATM/Tel1 functions in controlling resection, even if the resection machinery itself differs (e.g., role of EXO1).

### PRDM9-dependent modulation of local chromatin accessibility

How recombination machinery accesses a chromatinized donor is not well understood (Neale and Keeney 2006; Kobayashi et al. 2016). Average profiles around hotspots of H3K4me3 and H3K36me3 chromatin immunoprecipitation (ChIP) sequencing data show peaks of methylated nucleosome coverage flanking PRDM9 binding sites and the main cluster of DSB positions (**Figure 4A**) (Baker et al. 2014; Lange et al. 2016; Powers et al. 2016; Yamada et al. 2017). An implication is that the strand exchange machinery frequently invades segments of the homologous donor DNA that contain these methylated nucleosomes (if PRDM9 acted on the donor as well as the broken chromosome) plus additional flanking nucleosomes further away (**Figure 4B**). To more precisely define the chromatin structure at such recombination sites, we examined maps of bulk nucleosomes (ChIP input samples) liberated by micrococcal nuclease (MNase) from flow-sorted spermatocyte nuclei from B6 mice (Lam et al. 2019).

**Figure 4.**
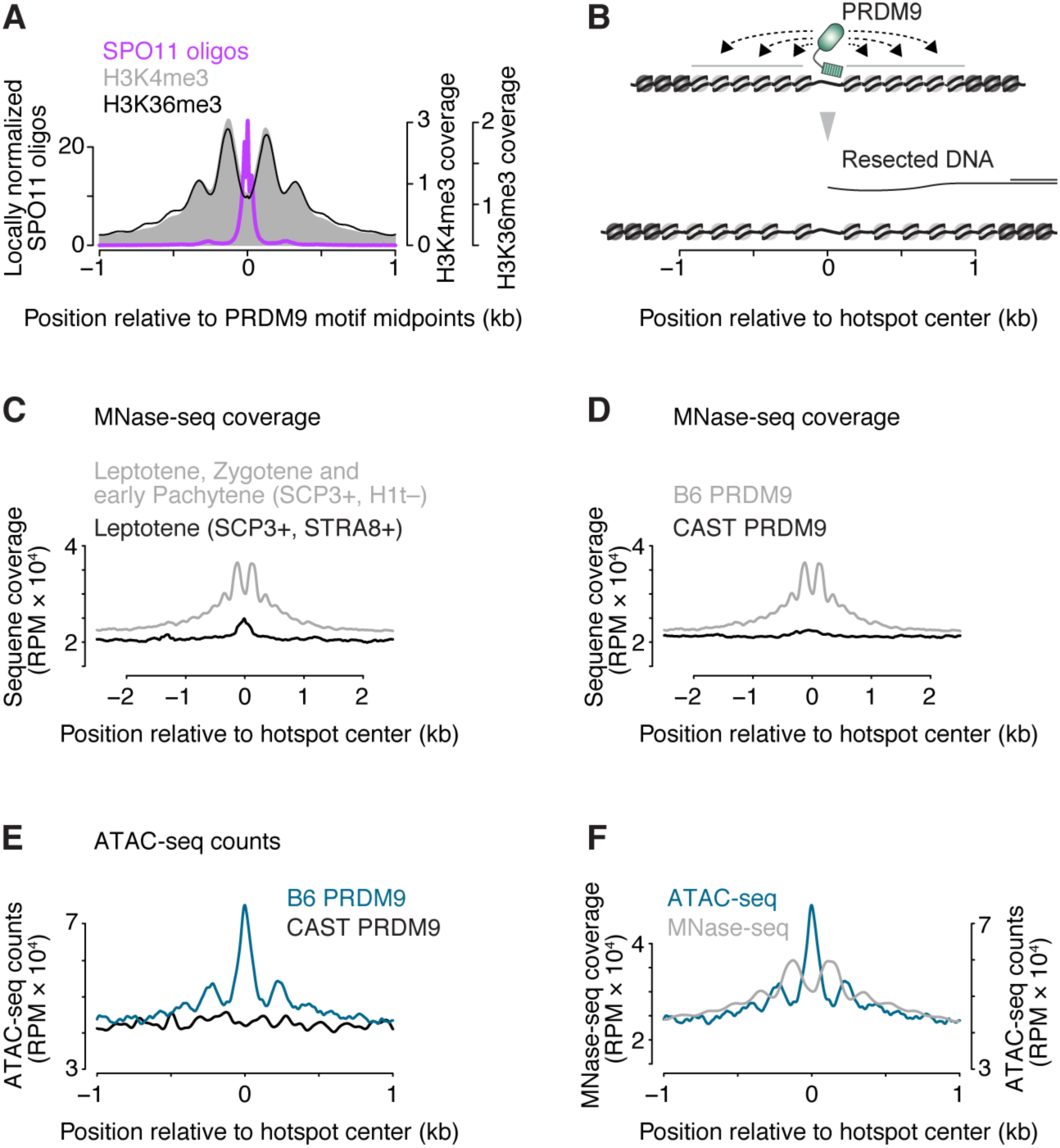
Increased chromatin accessibility at sites of PRDM9 activity. (A) Profiles of nucleosomes trimethylated at H3K4 or H3K36 around hotspot-enriched 12-bp PRDM9 motifs. Modified from (Yamada et al. 2017); methylated histone ChIP-seq data are from (Baker et al. 2014; Powers et al. 2016). (B) To-scale schematic illustrating how resected DNA of the broken chromosome compares to the zone over which PRDM9 influences chromatin structure. (C,D) MNase-seq coverage around hotspots. Shown are average profiles of sequence coverage from ChIP-seq input samples (before immunoprecipitation) prepared by MNase digestion of flow-sorted spermatocyte nuclei (Lam et al. 2019). Note that this represents total DNA liberated by MNase digestion, not just methylated nucleosomes. Panel C compares samples of the indicated stages of meiotic prophase I around DSB hotspots active in this strain (B6). Panel D compares the profile around B6 hotspots with the profile around hotspots not active in these strains (i.e., sites that can be targeted by the CAST version of PRDM9). (E) ATAC-seq cleavage profiles around active (PRDM9^B6^) and inactive (PRDM9^CAST^) hotspots. (F) Comparison of ATAC-seq cleavage positions with MNase-seq coverage maps. Signal was smoothed with a 51-bp Hann filter in **A**, and **C**-**F**.

MNase-seq coverage from cells in leptonema through early pachynema of the first meiotic prophase displayed a series of peaks flanking PRDM9 sites and extending 1 kb (∼6 nucleosomes’ worth) on either side on average (**Figure 4C**). At these stages, PRDM9-dependent methylation is present, DSBs are made, and recombination is occurring (Mahadevaiah et al. 2001; Moens et al. 2002; Guillon et al. 2005; Lam et al. 2019). A sample enriched for leptonema (i.e., early cells in which DSBs are just beginning to form) showed little indication of this signature (**Figure 4C**), which was also specific for sites of PRDM9 action (**Figure 4D**).

Importantly, we noted that this signature represents extra sequence coverage above the local baseline. If nucleosomes were becoming well positioned where they were already present but randomly distributed in the population, we would instead expect the peaks and valleys to oscillate above and below the baseline, i.e., the area under the coverage curve would have been unchanged. The increased MNase-seq coverage we observed instead could mean that PRDM9-modified sites have higher nucleosome occupancy, or, conversely, that greater chromatin accessibility allows MNase to more readily release nucleosomes.

To distinguish between these possibilities, we assayed testis cell chromtain structure in B6 mice using ATAC-seq, which assesses how accessible DNA is to Tn5 transposase (Buenrostro et al. 2013). We observed ATAC-seq read depth that was elevated above local baseline specifically around PRDM9 sites (**Figure 4E**). Tn5 integration positions—enriched in the central NDR and the linkers between flanking nucleosomes—were anticorrelated with MNase-seq coverage, as expected (**Figure 4F**).

We conclude that PRDM9 action causes changes in local chromatin structure that include a tendency toward positioning of nucleosomes but also greater overall accessibility. An earlier study also argued in favor of this idea, but without supporting evidence (Baker et al. 2014). Because neither the MNase-seq nor ATAC-seq maps should have been able to pick up a nucleosomal sequencing signature on the ssDNA of resected DSB ends, we infer that the increased chromatin accessibility is occurring on intact chromosomes.

## Discussion

Our results uncover both conserved and unique features of mouse meiotic resection. A surprisingly unique aspect was the minimal contribution of EXO1 to overall resection lengths, unlike in yeast. One possibility is that mouse meiocytes, similar to somatic cells, use additional resection activities that act redundantly with EXO1, such as DNA2 plus BLM or WRN helicase (Symington 2014).

Conserved features include a similar overall length scale for resection lengths in yeast and mouse despite large differences in genome size, and an important role for DMC1 in limiting the extent of resection. It is likely that assembly of DMC1 and/or RAD51 nucleoprotein filaments limits access of DSB ends to the resection machinery (Shinohara et al. 1997; Henry et al. 2006). In this context, the correlation of hyper-resection with absence of intermolecular recombination intermediates in *Atm*^*–/–*^ (but not *Spo11*^*+/–*^ *Atm*^*–/–*^) mutants points to defects in strand-exchange protein assembly and/or function affecting both resection and strand exchange.

Even more striking is the conservation of roles of ATM (Tel1) in controlling resection. ATM promotes resection in some contexts in somatic cells as well, although the mechanism is not well understood (e.g., Lee et al. 2018). Our findings provide the insight that ATM can influence resection in at least two ways. First, accumulation of unresected DSBs in *Atm*^*–/–*^ mutants suggests that ATM regulates MRE11 endonuclease activity, perhaps via phosphorylation of CtIP (Cartagena-Lirola et al. 2006). Second, the shorter resection tracts in *Atm*^*–/–*^ mutants indicate that ATM controls the extension of resection once it has begun. One possibility is that ATM directly stimulates the resection machinery and/or down-regulates inhibitors of resection. A non-exclusive possibility is that ATM recruits chromatin remodeling activities to DSBs (Chakraborty et al. 2018).

A striking feature of S1-seq maps is the hotspot-proximal signal that likely reflects intermolecular recombination intermediates. If these are D-loops, the spatial disposition of the signal suggests that they consist of invasion of an internal segment of the ssDNA and usually do not extend all the way to the 3′ end (**Figure S2C and S3**). A simple way to account for this would be if SPO11-oligo complexes cap the 3′ ends of resected DSBs, as we previously proposed (Neale et al. 2005) (**Figure S3**). Potentially consistent with such a cap being biochemically stable, recombinant yeast Spo11 complexes bind very tightly (even without a covalent end-linkage) to DNA ends *in vitro* (Claeys Bouuaert and Keeney, in preparation). Moreover, the exonucleolytic END-seq method (Canela et al. 2019) detects a sequencing signal in mouse testis cells that resembles unresected DSBs, which would be predicted for a SPO11-oligo-capped end (A. Nussenzweig, personal communication, and ref. (Mahgoub et al. 2019)).

If correct, an intriguing implication is that capping would provide a mechanism to maintain the strand-exchange intermediate in a poised state that retains potential to move either forward or backward. Displacing the SPO11-oligo complex (Neale et al. 2005) or removing the uninvaded 3′ end by cleaving the flap (Peterson et al. 2019) would free a 3′-OH end to prime DNA synthesis and drive D-loop extension. Conversely, reversal of strand exchange would leave the broken end with a ssDNA gap that could be filled in by DNA synthesis primed from the 3′ end of the SPO11 oligo, potentially allowing non-homologous end joining as a backup repair pathway.

Finally, our studies uncover a previously undocumented remodeling of chromatin structure at sites of PRDM9 action. The more open chromatin structure observed with both MNase digestion and ATAC-seq may reflect wider linkers, displacement of linker histones, less tightly bound nucleosomal particles, and or dynamic nucleosome removal and redeposition such that a fraction of DNA molecules is free at steady state. Regardless of the source, this open chromatin structure seems likely to be important for SPO11 access. Moreover, a further key implication is that PRDM9 action on the uncut homolog generates a more accessible chromatin structure precisely where its broken partner needs to engage it for repair (**Figure 4B**). Thus, this greater accessibility may provide a mechanistic explanation for observed repair differences between hotspots that are symmetrically vs. asymmetrically bound by PRDM9 (Davies et al. 2016; Hinch et al. 2019; Li et al. 2019).

## Materials and Methods

### Mice

Experiments conformed to the US Office of Laboratory Animal Welfare regulatory standards and were approved by the Memorial Sloan Kettering Cancer Center Institutional Animal Care and Use Committee. Mice were maintained on regular rodent chow with continuous access to food and water until euthanasia by CO_2_ asphyxiation prior to tissue harvest. Previously described *Spo11* (Baudat et al. 2000), *Atm* (Barlow et al. 1996), and *Exo1*^*DA*^ (Zhao et al. 2018) mutations were maintained on a congenic B6 strain background. The *Dmc1* mutation (Pittman et al. 1998) was maintained on a mixed (129/SV and B6) background. B6xPWD F1 hybrid mice (semi-fertile) were generated by crossing B6 (stock number 000664) female mice and PWD/PhJ (stock number 004660) male mice obtained from The Jackson Laboratory.

### S1-seq

#### Testis dissociation

Unless otherwise stated, testis cells from12-dpp juvenile mice were obtained as described previously (Cole et al. 2014). Briefly, testes were decapsulated, and incubated in Gey’s balanced salt solution (GBSS, Sigma) with 0.5 mg/ml collagenase type 1 (Worthington) at 33 °C for 15 min. Seminiferous tubules were then rinsed three times and further treated with 0.5 mg/ml trypsin (Worthington) and 1 µg/ml DNase I (Sigma) at 33 °C for 15 min. Trypsin was inactivated with 5% FCS and tubules were further dissociated by repeated pipetting. Cells were passed through a 70-µm cell strainer (BD Falcon) and washed 3 times in GBSS.

#### DNA extraction in plugs

Cells were embedded in plugs of 0.5% low-melting point agarose (Lonza) in GBSS. One plug mold (Bio-Rad) was used per 2 testes (1∼2 million cells per plug). Plugs were incubated with 50 µg/ml of proteinase K (Roche) in lysis buffer (0.5 M EDTA pH 8.0, 1 % N-lauroylsarcosine sodium salt) overnight. Plugs were washed 5 times with TE (10 mM Tris-HCl pH 7.5, 1 mM EDTA pH8.0), and then incubated with 50 µg/ml RNase A (ThermoFisher) at 37 °C for 3 hours. They were washed 5 times with TE and stored in TE at 4 °C

#### S1-seq library preparation

In-plug overhang removal with S1 and adaptor ligation were performed as previously described (Mimitou et al. 2017; Mimitou and Keeney 2018). For ExoT-seq, the S1 treatment step was replaced with the following: Plugs were washed in 1 ml of NEBuffer 4 for 15 min 3 times, treated with 75 units of exonuclease T in 100 µl of NEBuffer 4 at 24°C for 90 min and washed 3 times with 8 ml of TE for 15 min (Canela et al. 2019). After adaptor ligation, plugs were washed 3 times in TE and incubated in TE at 4 °C overnight to diffuse excessive unligated adaptors out of plugs. Agarose was then digested by the Epicentre GELase Enzyme Digestion protocol. DNA was fragmented by vortex and further sheared to DNA fragment sizes ranging between 200-500 bp with a Bioruptor^®^ waterbath sonicator (Diagenode) at 4 °C for 40 cycles of 30 seconds ON/OFF at the middle power setting. DNA was purified by ethanol precipitation and dissolved in 100 µl TE. Fragments containing the biotinylated adaptor were purified with streptavidin, ligated to adaptors at sheared end and amplified by PCR as previously described (Mimitou et al. 2017; Mimitou and Keeney 2018). PCR products were purified with 0.9× AMPure XP Beads (Beckman Coulter) to remove primer dimers. DNA was sequenced on the Illumina HiSeq platform in the Integrated Genomics Operation at Memorial Sloan Kettering Cancer Center.

#### Mapping and pre-processing

Sequence reads were mapped onto the mouse reference genome (mm10) by bowtie2 version 2.2.1 (Langmead et al. 2009) with the argument -X 1000. Uniquely and properly mapped reads were counted at which a nucleotide next to biotinylated adaptor DNA was mapped (this corresponds to a position of a resection endpoint).

### ATAC-seq

Testis cells were isolated from 12-dpp juvenile mice as described for S1-seq sample preparation. Subsequent ATAC-seq library preparation and sequencing were performed in the Integrated Genomics Operation at Memorial Sloan Kettering Cancer Center as described (Buenrostro et al. 2013) with slight modifications. Briefly, nuclei extracted from 50,000 testis cells were treated with Tn5 transposase at 42 °C for 45 min. Libraries were amplified and sequenced on the Illumina HiSeq platform. Reads were mapped as described for S1-seq data processing. Uniquely and properly mapped reads were counted at which Tn5 insertion positions were mapped.

### Other datasets and data availability

Raw and processed sequencing S1-seq and ATAC-seq data were deposited at the Gene Expression Omnibus (GEO) (accession number GSE141850). We used SPO11-oligo, SSDS and H3K4me3 data from GEO accession numbers GSE84689, GSE35498, and GSE52628, respectively (Brick et al. 2012; Baker et al. 2014; Lange et al. 2016). For SPO11-oligo maps, we used either “B6” or “*Atm* null 1” dataset, which were respectively from wild-type mice from a pure B6 background, or *Atm*^*–/–*^ mice from a mix of B6 and 129/Sv strain backgrounds, which carry the same *Prdm9* allele. Unless otherwise stated, hotspots analyzed in this study were the 13,960 hotspots previously identified using uniquely mapped SPO11-oligo reads (Lange et al. 2016). The hotspot center was defined as the position of the smoothed peak in the SPO11-oligo density.

### Quantification and statistical analyses

Statistical analyses were performed using R versions 3.2.3 to 3.3.1 (http://www.r-project.org) (R Core Team 2018). Statistical parameters and tests are reported in the figures and legends.

## End Matter

### Author Contributions and Notes

S.Y. performed experiments. S.Y. analyzed S1-seq data with contributions from A.G.H. K.H. performed an initial ATAC-seq analysis. Y.Z. and W.E. generated *Exo1*^*DA*^ mice. S.Y. and S.K. designed the study and wrote the paper with contributions from A.G.H.

The authors declare no conflict of interest.

## Acknowledgments

We thank A. Viale, N. Mohibullah, and R. Patel (MSKCC Integrated Genomics Operation) for sequencing and ATAC-seq library preparation; E. Mimitou for advice during adaptation of the S1-seq method; S. Peterson and M. Jasin (MSKCC), P. Donnelly (University of Oxford), and A. Nussenzweig, W. Wu, and J. Paiano (NIH) for discussions and sharing unpublished data; P.C. Huang (MSKCC) for ExoT treatment optimization; P.C. Huang and H. Murakami (MSKCC) for help in analysis of recombination intermediates; and members of the Keeney and Jasin labs for discussions. MSKCC core facilities were supported by NIH Cancer Center Core Grant P30 CA008748. This work was supported by NIH grant R35 GM118092 (to S.K.).

---

**Figure S1.**
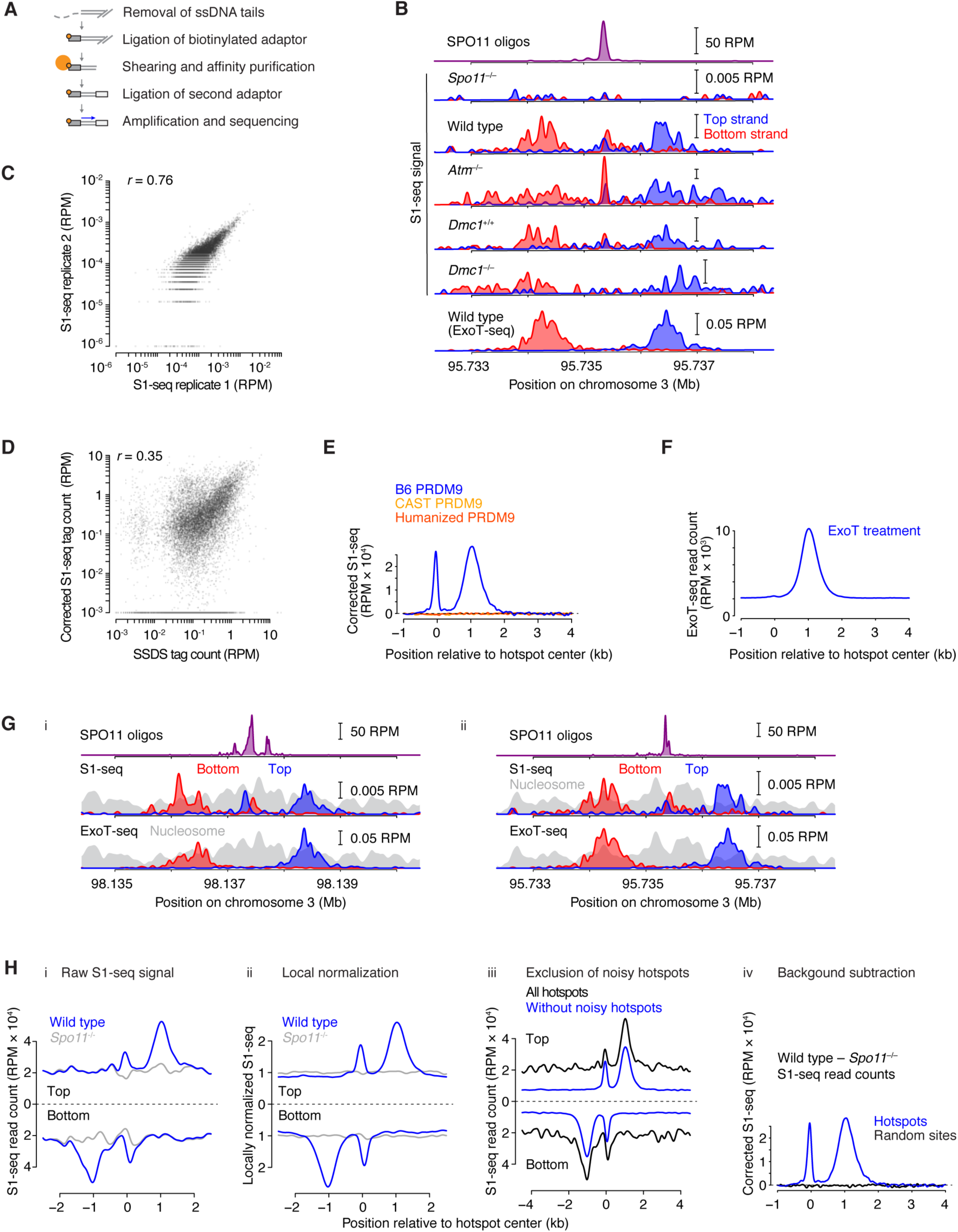
S1-seq maps in mouse spermatogenesis. (A) Schematic of steps in S1-seq library preparation. After removal of ssDNA ends with S1 nuclease, biotinylated adaptors are ligated to the resulting blunt DNA ends. The genomic DNA is then sheared by sonication and the biotinylated ends are affinity-purified using streptavidin. Ligation of a second adaptor to the sheared end allows PCR amplification and deep sequencing. (B) S1-seq at a second representative DSB hotspot, presented as for **Figure 1B**. (C) Reproducibility (Pearson’s *r*) of S1-seq maps. S1-seq read counts from biological replicates from wild-type mice were summed within a 5,001-bp window around each hotspot center. Each point represents one hotspot. Lower sequencing coverage in replicate 2 resulted in some weak hotspots showing no or few S1-seq tags. Noisy hotspots (n = 2,266), defined as as those with high read counts (0.00025 RPM within 5,001-bp window around hotspot centers) in biological-replicate-averaged *Spo11*^*–/–*^ data, were excluded from the analysis. Hotspots with no S1-seq tags (n = 57) were not used for Pearson’s *r* calculation but set as 10^−6^ for plotting purposes. (D) Correlation (Pearson’s *r*) of S1-seq read count (summed as in **Figures 1D and S1C**) with DSB intensity measured by SSDS (Brick et al. 2012). S1-seq signal was background-corrected by subtracting *Spo11*^*–/–*^ signal from the wild-type signal. Hotspots with ≤0 corrected S1-seq tag counts (n = 4,050) were excluded for Pearson’s *r* calculation. Hotspots with ≤10^−3^ corrected S1-seq tag counts (n = 4,071) were set as 10^−3^ for plotting purposes. (E) PRDM9-specificity of S1-seq signal. SPO11-specific S1-seq signal was averaged around hotspots in the B6 genome that can be targeted by the PRDM9 protein carried by CAST mice. The blue profile (from PRDM9^B6^-targeted hotspots) is reproduced from **Figure 1F** for comparison. (F) Average profile around hotspots for sequencing maps generated by digestion with the ssDNA-specific exonuclease ExoT instead of S1 endonuclease. Note the virtual absence of central sequencing signal (position 0 relative to hotspot centers). (G) No obvious correlation of S1-seq resection end points with nucleosome positions. Strand-specific S1-seq maps for the representative hotspots from **Figures 1B** (panel i) **and S1B** (panel ii) are compared with bulk MNase-seq coverage maps from SYCP3-positive, histone H1t-negative spermatocyte nuclei (mixed leptonema, zygonema, and early pachynema) (Lam et al. 2019). Peaks in the resection endpoint maps do not appear to systematically correspond to peaks or valleys in the MNase-seq maps. However, we note that there appears to be considerable heterogeneity in nucleosome positioning across the population of cells assayed, as indicated by the lack of clearly defined linker regions in the MNase-seq coverage. Therefore, it is likely that attempts to compare the population-average distributions of S1-seq and MNase-seq are uninformative about the relation of resection stopping points with underlying chromatin structure. Moreover, even in yeast such a spatial correlation is modest despite a clear causal relationship between chromatin structure and resection endpoints (Mimitou et al. 2017) and despite presence of well positioned nucleosome arrays (Jiang and Pugh 2009). We thus consider it likely that chromatin structure influences resection profiles in mouse as it does in yeast, but available population-average methods are not well suited to reveal this influence. (H) Comparison of different methods for background subtraction and normalization. i) Raw S1-seq signal. S1-seq signal around all hotspots (n = 13,960) was averaged for wild-type and *Spo11*^*–/–*^ B6 mice. ii) Local normalization. Data were locally normalized by dividing the signal at each base pair by the mean signal within each 2,501-bp window around hotspot centers, then were averaged across all hotspots. This local normalization weights each hotspot equally, without regard for DSB frequency, so it facilitates analysis of per-hotspot (rather than per-DSB) spatial patterns. iii) Exclusion of noisy hotspots. We defined noisy hotspots (n = 4,609) as those with high read counts (0.0002 RPM within 10,001-bp window around hotspot centers) in biological-replicate-averaged *Spo11*^*–/–*^ data. The mean S1-seq profiles with or without the noisy hotspots were averaged. iv) Background-subtracted (SPO11-dependent) signal. The *Spo11*^*–/–*^ S1-seq signal was subtracted from the wild type signal, then top and bottom strand reads were averaged after co-orienting by rotation around the hotspot centers. The graph shows a comparison of the SPO11-dependent S1-seq signal around all hotspots (reproduced from **Figure S1E**) or the same number of randomly chosen sites (n = 13,960). Note that the SPO11-dependent S1-seq signal was essentially zero at random sites, confirming specificity of this method. Signal was smoothed with a 151-bp Hann filter in **B, E**-**F, G**, and **H.iv**, and with a 401-bp Hann filter in **H.i**-**iii**.

**Figure S2.**
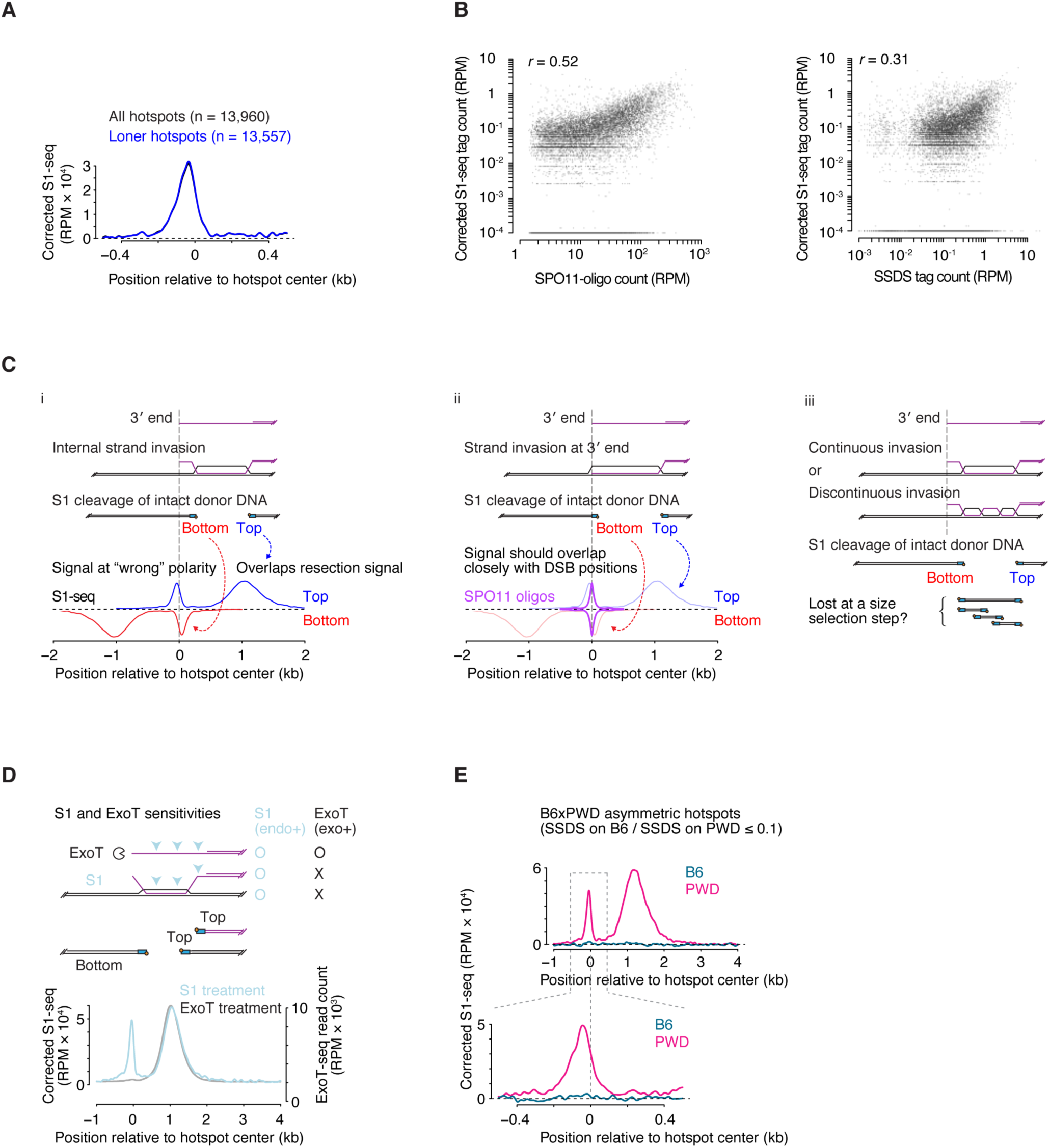
Intermolecular recombination intermediates. (A) Central S1-seq signal is not attributable to resection from adjacent hotspots. Central signal is compared for all hotspots and “loner” hotspots, i.e., those left after removing hotspots with another hotspot located within 2 kb. (B) Recombination intermediate S1-seq signal correlates with hotspot strength as measured by SPO11-oligo sequencing (left panel) or SSDS (right panel). Each point is a SPO11-oligo hotspot, with S1-seq signal summed from –100 to +300 bp (bottom strand) and –300 to +100 bp (top strand) around hotspot centers. S1-seq signal was background-corrected by subtracting *Spo11*^*–/–*^ signal from the wild-type signal. Hotspots with ≤0 corrected S1-seq tag counts (n = 5,247) were excluded for Pearson’s *r* calculation. Hotspots with ≤10^−4^ corrected S1-seq tag counts (n = 5,264) were set as 10^−4^ for plotting purposes. (C) Additional schematics illustrating features of the recombination intermediate signal. (i) Internal invasion with uninvaded 3′ end, reproduced from **Figure 2D** to facilitate comparison. (ii) Alternative scenario with fully invaded 3′ end (canonical model (Hunter 2015)). Note that this scenario predicts that the central signal should overlap closely with the positions of DSBs, which is not observed. (iii) Internal invasions could involve a single continuous D-loop, or multiple smaller invasions. In either case, S1 cleavage fragments in between the most proximal and most distal positions would not yield sequenceable library fragments because these would be eliminated by the fractionation step that selects for high molecular weight DNA after adaptor ligation (Methods). (D) Comparison of predicted sensitivities of resected DSBs and D-loops to S1 vs. ExoT nucleases. Average sequencing profiles are reproduced from **Figures 1F and S1F**. (E) S1-seq in B6xPWD F1 hybrid mice. S1-seq signal was background-corrected with signal from F1 hybrid mice at 9 dpp, when few meiotic cells have yet formed. We attempted to test if the central signal reflects invasion of the homolog by generating S1-seq maps from F1 hybrid mice in which DSBs at certain hotspots occur only or predominantly on one of the parental chromosomes because of strain-of-origin differences in PRDM9 binding (Davies et al. 2016; Smagulova et al. 2016). At highly asymmetric hotspots where PRDM9^B6^ preferentially targets the PWD chromosomes (defined as those where the fraction of total SSDS reads that derived from the B6 chromosome was ≤0.1; n = 1,220), we observed a central signal derived only from the same chromosomes that were undergoing DSB formation, i.e., the PWD chromosomes. (Note that SPO11-oligo maps are only available for PRDM9^B6^-targeted DSBs. Because detecting the central signal requires the high spatial precision afforded by SPO11-oligo mapping of hotspot centers, we cannot examine PRDM9^PWD^-targeted hotspots that are as yet only defined using SSDS. Moreover, it was not informative to examine the reciprocally asymmetric hotspots where PRDM9^B6^ targets the B6 genome, because these are very weak hotspots due to hotspot erosion from biased gene conversion.) This result suggests that at least some of the putative recombination intermediates giving rise to the central S1-seq signal involve invasion of the sister chromatid. One possibility is that all of the central signal reflects inter-sister joint molecules, in which case these may be related to a proposed “quiescent-end” complex that inhibits recombination by one end of a DSB while the other end is released to search for (non-sister) homology (Storlazzi et al. 2010). An alternative, however, is that interhomolog joint molecules contribute to the signal at symmetric hotspots (e.g., in B6 mice) but are not detectable by S1-seq at asymmetric hotspots. This could be because steady-state levels are reduced because of more dynamic turnover or a delay in formation, and/or because D-loop boundaries become more variable and thus too blurred to detected in population average. Both effects (reduced levels and altered distributions) can easily be envisioned to accompany a lack of PRDM9-dependent opening of chromatin structure on the intact homolog. Regardless of the cause, however, if this hypothesis is correct, then the same polymorphisms that are required to discriminate intersister from interhomolog recombination prevent the interhomolog events from being detected, which would mean that available methods are unable to define the chromosome of origin. Signal was smoothed with a 151-bp Hann filter in **D** (lower panel), and **E** (upper panel), and with a 51-bp Hann filter in **A**, and **E** (lower panel).

**Figure S3.**
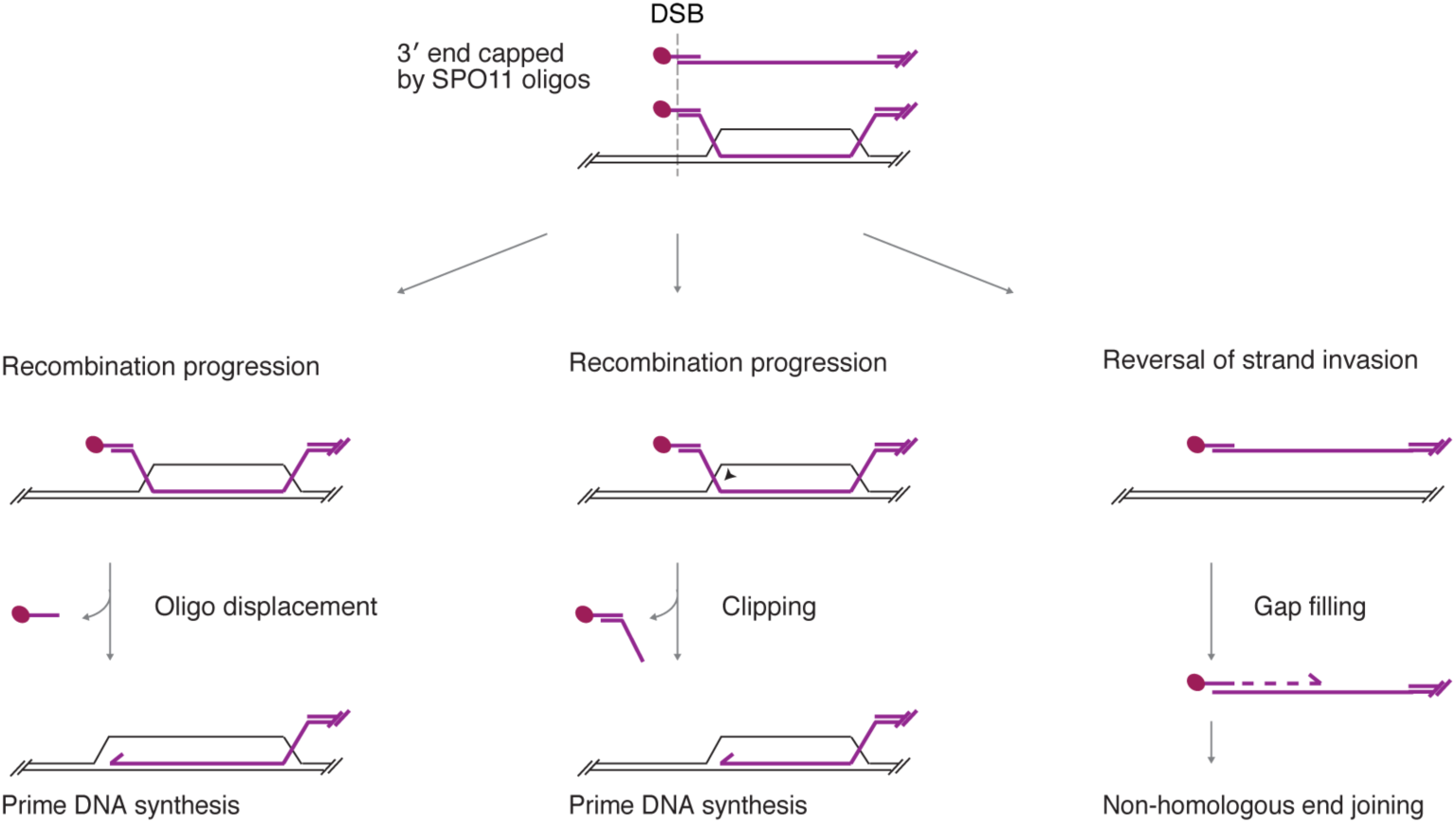
Model for capping of the 3′ ends of DSBs by annealed SPO11-oligo complexes. Such caps, originally proposed in part because SPO11 oligos are long enough to remain base paired at physiological temperature (Neale et al. 2005), might explain the lack of strand invasion by the 3′ end and would be predicted to inhibit extension of the invading strand by DNA polymerase. Such a structure might be poised between moving forward into the recombination reaction (by removal of the SPO11-oligo complex or clipping of the uninvaded flap) or reversal and repair by nonhomologous end joining as a backup pathway after gap fill-in primed from the 3′ end of the SPO11 oligo.

